# Predator decision-making shapes the dynamics and stability of mimicry systems

**DOI:** 10.1101/2025.10.15.670298

**Authors:** Yi Sun, Po-Ju Ke

## Abstract

Mimicry is an anti-predator strategy in which prey species (the mimic) resemble an unprofitable species (the model) to deceive predators. Despite theoretical expectations for perfect mimicry, imperfect mimicry, where the mimic resembles its model imperfectly, is widespread in nature. To understand how imperfect mimicry can persist ecologically, we studied the effect of different predator recognition processes on the dynamics and stability of various mimicry systems. Specifically, we extended a dynamical model that integrates optimal foraging and signal detection theories by introducing a novel abundance-dependent recognition mechanism, where predators’ perception of the similarity between mimic and model is influenced by the relative abundance of prey types. We demonstrate that intermediate similarity promotes stable community dynamics and increases mimic abundance in single Batesian mimicry systems. Moreover, abundance-dependent recognition leads predators to reduce attack on mimics with low morphological similarity, further contributing to system stability. Extending the framework to a multi-mimicry system, we find that Batesian and Müllerian mimics have contrasting effects: intermediate Batesian similarity continues to stabilize the system, while high Müllerian similarity provides additional protection and can offset destabilization caused by highly similar Batesian mimics. Our study offers a novel explanation for the prevalence of imperfect mimicry in nature and highlights how recognition processes shape the ecological stability of mimicry systems.

## Introduction

Mimicry is a complex anti-predator adaptation that involves interactions among three main players: the mimic, the model, and the predator (Ruxton *et al*., 2004, Quicke, 2017). In this system, the model is an unprofitable prey that displays aposematic signals to advertise its unprofitability. The mimic, on the other hand, displays phenotypes that resemble those of the model, including behavior, chemical compounds, and, more commonly, morphological characteristics such as color, pattern, and shape (Kikuchi *et al*., 2013). These phenotypes deceive predators into misidentifying the mimic as the unprofitable model. One can categorize a mimicry system based on the mimic’s profitability to the predator, with Batesian and Müllerian mimicry being the two most widely recognized forms. In Batesian mimicry, the mimic is a profitable prey that reduces its predation risk by resembling an unprofitable model. However, the model in this system involuntarily suffers higher mortality as the presence of the Batesian mimic confuses the predators, thereby reducing the effectiveness of the model’s aposematic display (Bates, 1862). In Müllerian mimicry, the mimic is an unprofitable prey that shares a common aposematic signal with the model. Unlike Batesian mimicry, both the mimic and the model benefit from their resemblance in Müllerian mimicry due to the positive reinforcement of the aposematic signal (Müller, 1879, Vane-wright, 1980, Mallet & Joron, 1999, Ruxton *et al*., 2004). It has been suggested that natural selection should favor mimics that closely resemble their model in both mimicry systems, resulting in highly similar mimic–model pairs (Nur, 1970, Sherratt, 2002, Ruxton *et al*., 2018). However, empirical evidence shows that many mimics do not closely resemble their models despite the apparent advantage of high similarity, a phenomenon known as imperfect mimicry (Sherratt, 2002, Kikuchi *et al*., 2013, Sherratt & Peet-Paré, 2017, Bosque *et al*., 2018, McLean *et al*., 2019).

This contradictory phenomenon raises the question: what mechanisms allow imperfect mimicry to persist in nature? Many hypotheses and theories have been proposed to explain the existence of imperfect mimicry (Penney *et al*., 2012, Pfennig & Kikuchi, 2012, Kikuchi *et al*., 2013). On one hand, evolutionary explanations, such as the “chase-away” hypothesis, predict that imperfect mimicry exists because models evolve away from their mimic (Nur, 1970, Oaten *et al*., 1975, McGuire *et al*., 2006, Franks *et al*., 2009, Akcali *et al*., 2018) (also see Sherratt (2002), Penney *et al*. (2012), Johnstone (2002), Pfennig & Mullen (2010), Tomizuka & Tachiki (2024) for more evolutionary hypotheses). On the other hand, ecological explanations, such as the “eye of the beholder” hypothesis, argue that what is perceived as imperfect mimicry by humans may in fact be effective from the perspective of the predator (Cuthill & Bennett, 1993, Dittrich *et al*., 1993). Furthermore, the “multiple models and multiple predators” hypothesis suggests that imperfect mimicry could arise either because the mimic resembles multiple models or because multiple predators rely on different signals to detect the mimic (Edmunds, 2000, Sherratt, 2002, Pekár *et al*., 2011). As a result, the mimic might resemble many aposematic signals from different models, resulting in a signal that partially resembles several model species but does not closely match any single one (also see Kikuchi *et al*., 2013 for additional ecological hypotheses).

In addition to morphological similarities, different types of predator recognition processes could also influence the dynamics of mimicry systems (Darst, 2006, Chittka & Osorio, 2007). These recognition processes shape the realized similarity perceived by the predator, which may differ from the actual mimic–model morphological similarity. Predator recognition can be affected by the learning efficiency of the predator (Huheey, 1964) or by the abundance of different prey items (Nelson *et al*., 2010). Müllerian mimicry is a classic example of such abundance-dependent recognition — a higher abundance of unprofitable mimics increases predator encounters and reinforces the association between the aposematic signal and unprofitability (Müller, 1879). This type of abundance-dependent recognition is particularly important on an ecological timescale as it creates intricate feedback between prey abundance, predator recognition, predator attack decisions, and subsequent community dynamics.

Despite numerous hypotheses being proposed to address the emergence of imperfect mimicry, its effects on population dynamics and the stability of mimicry systems remain unclear. Understanding these ecological mechanisms and their consequences clarifies how imperfect mimicry persists over short ecological timescales, which is essential for setting the stage for its evolution over longer timescales. Previous theoretical studies addressing the stability of mimicry systems often lacked comprehensive dynamics of the entire system: some overlooked the dynamics of the predator (e.g., Getty, 1985), while others omitted both the predator and the non-mimetic alternative prey (e.g., Yamauchi, 1993, Kumazawa *et al*., 2006). Other studies have focused on how mimicry dynamics are shaped by factors such as the influx of alternative prey, the degree of similarity between the mimic and the model, and the level of defense equipped by the model species (Brower & Moffitt, 1974, Rowell-Rahier *et al*., 1995, Kikuchi *et al*., 2022). Many of these theoretical studies employed optimal foraging theory, which assumes that predators make attack decisions based on properties of the prey items (i.e., their profitability and abundance; Charnov, 1973, 1976, Stephens & Krebs, 1986).

Although previous theoretical studies on mimicry population dynamics have mostly focused on single Batesian mimic system based on morphological similarity, natural mimicry often forms multi-mimicry complexes involving multiple types of mimicries and various recognition processes. Specifically, multi-mimicry complexes involving both Batesian mimicry and Müllerian mimicry are common in nature (e.g., *Heliconius* butterfly Quicke, 2017 and *Pachyrhynchus* weevils Schultze, 1923), where abundance-dependent recognition may also play an important role. With a greater number of prey types involved, the predator needs to juggle between multiple mimic–model similarities and prey profitabilities when making attack decisions. Importantly, Müllerian mimics differ fundamentally from Batesian mimics: the resemblance between the Müllerian mimic and the model produces a positive reinforcement that provides both species greater protection, whereas the resemblance between the Batesian mimic and the model imposes a cost on the model species. Therefore, Batesian and Müllerian mimics may differentially influence the dynamics of the multi-mimicry system. To investigate the persistence of imperfect mimicry in natural systems with both Batesian mimicry and Müllerian mimicry, it is essential to consider the interactive effect of their similarity with the model and their profitability for the predator.

Here, we extended the framework proposed by Kikuchi *et al*. (2022), which offers an exciting opportunity to investigate the persistence of imperfect mimicry and its resulting community dynamics by integrating optimal foraging theory into a dynamical model of the full mimicry system. By exploring the impact of mimic–model similarity in a single Batesian mimicry system, we offer a novel perspective on the ecological dynamics of mimicry systems: imperfect mimicry can stabilize community dynamics and lead to a higher mimic abundance. We then explore how this stability pattern conferred by imperfect mimicry varied with different predator recognition mechanisms. To this end, we formulated a novel phenomenological representation of the predator’s recognition process, which depends on both the innate mimic–model similarity (i.e., morphological similarity based on prey traits) and the abundance ratio of the mimic and the model. Finally, we expanded the framework to a multi-mimicry system encompassing both Batesian and Müllerian mimicry. We explored how the dynamics of the multi-mimicry system are influenced by the similarities between different mimics and the model under the aforementioned abundance-dependent recognition-determining process. Overall, we show that imperfect mimicry can, counterintuitively, promote higher mimic abundance and greater stability across a range of mimicry systems that differ in predator recognition mechanism and the number of mimic species involved.

## Method

We used a theoretical model to study how mimic–model similarity and different predator recognition processes influence predation decisions and the dynamics of mimicry systems. Our ordinary differential equation (ODE) model is built upon the theoretical framework of Kikuchi *et al*. (2022), which combines optimal foraging theory and signal detection theory to simulate the dynamics of mimicry systems. We extended this framework to investigate how mimic–model similarity and predators’ abundance-dependent recognition influence the abundance of mimic species and the stability of the system through predator decision-making (Fig. 1). In the following sections, we first introduce the foundations determining the predator decision-making process, including optimal foraging theory, signal detection theory, and abundance-dependent recognition. We then present the population dynamic framework that governs the abundance of species within the mimicry system. Finally, we describe the numerical setup used in our study.

**Figure 1:**
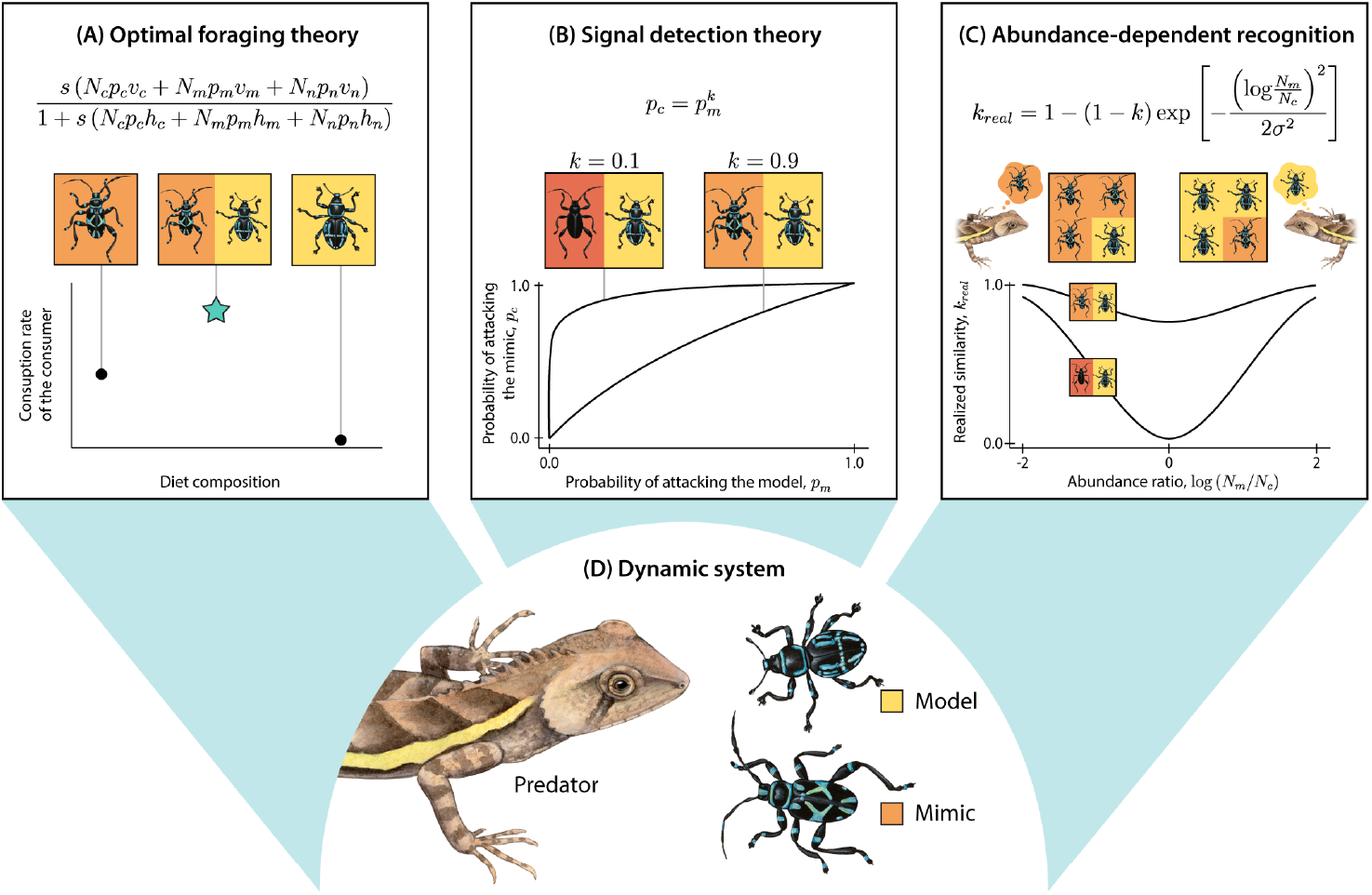
Illustrative figure demonstrating how our study incorporates (A) optimal foraging theory (i.e., predator optimizing attack probability *p*_*i*_ to maximize foraging gain), (B) signal detection theory (i.e., non-independence of attack probabilities via mimic–model morphological similarity *k*), and (C) abundance-dependent recognition (i.e., predator realized recognition *k*_*real*_ also dependent of model–mimic abundance ratio) into a (D) dynamic system of mimic–model–predator. Here, only a single Batesian mimic is shown for simplicity (so the secondary subscript *j* in 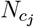 is omitted). In panels (B) and (C), we depict two scenarios with low (*k* = 0.1; red mimic) and high (*k* = 0.9; orange mimic) innate mimic–model morphological similarity *k*. See the main text for the mathematical description of the full multi-mimic model.

### Predator decision-making process

#### Optimal foraging theory

We followed the classic assumption that predators determine their probability (*p*_*i*_) of attacking three different types of prey items – mimics (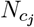; *j* = 1, 2, *· · ·, n*), model (*N*_*m*_), and alternative prey (*N*_*n*_) – through optimal foraging theory (Charnov, 1973, 1976; Fig. 1A). Assuming a multi-prey Holling type-II functional response, the term that predators attempt to optimize while making foraging decisions is:

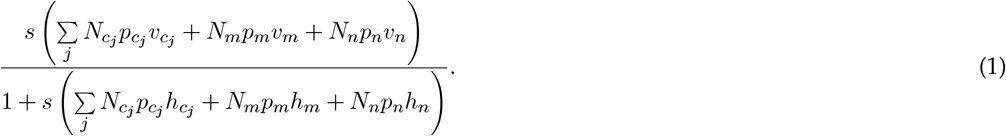

Here, *p*_*i*_ represents the probability of the predator attacking prey item *i* (*i* = *c*_*j*_, *m, n* for mimics, model, and alternative prey, respectively). The parameter *v*_*i*_ represents the value that the predator gains from consuming one individual of prey item *i*, whereas *h*_*i*_ represents the time predators spend handling one individual of prey item *i*. Therefore, each *N*_*i*_*p*_*i*_*v*_*i*_ term in the numerator represents the value gain of consuming species *i*, whereas each *N*_*i*_*p*_*i*_*h*_*i*_ term in the denominator represents the cost of consuming species *i*. The parameter *s* represents the predator’s search rate, which is assumed to be equal for all prey items. Here, we defined a prey item’s profitability based on its 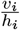 (Charnov, 1976). We further assumed that at least one mimic species is a Batesian mimic and will have the highest profitability, followed by the alternative prey, and, finally, the model. Therefore, when there is only one mimic species (i.e., the Batesian mimic; *j* = 1), this parameterization ensures that 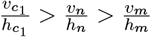, i.e., the (Batesian) mimic will be the most profitable prey. In classic optimal foraging theory, which assumes that predators have complete information about the system and can instantaneously recognize different prey types, predator behavior always follows an all-or-nothing manner (Charnov, 1976, Stephens & Krebs, 1986). That is, predators will either always include that prey in their diet (*p*_*i*_ = 1) or never include that prey in their diet (*p*_*i*_ = 0) upon encounter, with a decision-switching threshold determined by the profitability and abundance of prey items (Stephens & Krebs, 1986).

#### Signal detection theory

In mimicry systems, however, predators may experience difficulty in distinguishing between the mimic and the model. Such imperfect information results in a non-binary consumption probability due to the numerical connection between 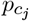 and *p*_*m*_. Mathematically, this means that the predator cannot make independent decisions on 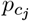 and *p*_*m*_ to optimize eq. 1; (Getty, 1985, Kikuchi *et al*., 2022). One can first assume that the predator has a fixed recognition ability, which, in the simplest case, depends solely on the morphological similarity between the mimic and the model. Following Getty (1985), the probabilities of attacking the mimic and the model are linked by a power law function:

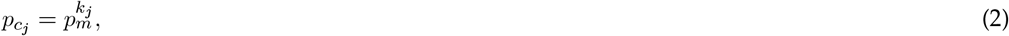

with the exponent *k*_*j*_ *∈* [0, 1] representing the morphological similarity between mimic 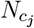 and the model *N*_*m*_ (Fig. 1B). When *k*_*j*_ = 0, the predator can distinguish between the mimic and the model perfectly, indicating complete discriminability from the predator’s perspective. Following classic optimal foraging, in a single Batesian mimic system (i.e., *j* = 1), the predator will always attack the mimic (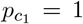) due to its higher profitability (i.e., 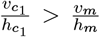). On the other end of the spectrum, the two prey items are morphologically identical from the predator’s perspective when *k*_*j*_ = 1 (i.e., perfect mimicry); the predator would treat the mimic and the model as the same species and either attack or reject them altogether (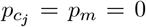 or 1). However, in most natural cases, mimicry falls between the two extreme cases, and *k*_*j*_ will be a number between 0 and 1. Here, 0 *< k*_*j*_ *<* 1 represents the scenario when imperfect information interferes with optimal foraging decision-making, leading to non-binary optimal foraging decisions (i.e., 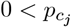, *p*_*m*_ *<* 1). As *k*_*j*_ approaches one, predators experience greater difficulty distinguishing the mimic from the model, leading to a higher likelihood of attacking the wrong prey (i.e., a higher perceived mimic–model similarity). For our purpose, we defined intermediate similarity as 0.5 *< k*_*j*_ *<* 0.8. At the same time, the attack probability of the alternative prey remains consistent with the classic optimal foraging theory (i.e., *p*_*n*_ = 0 or 1).

#### Abundance-dependent predator recognition

In addition to the morphological similarity between the mimic and the model, we considered a novel predator recognition process that determines the predator’s realized similarity *k*_*real, j*_ based on three factors. First, the realized similarity is constrained by the innate morphological similarity between mimics and the model, which corresponds to *k*_*j*_ in eq. 2 and represents the lowest value (highest distinguishability) that *k*_*real, j*_ can achieve. Second, the realized similarity is influenced by the abundance ratio of two prey categories: profitable (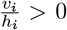) and unprofitable 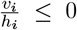) prey. When unprofitable prey is significantly more abundant than profitable prey, predators interacting with this mimicry complex are more likely to encounter unprofitable prey. This causes the predator to associate the shared morphological traits with the unprofitable prey, thereby perceiving both prey types as unprofitable prey. The opposite outcome occurs when the abundance of the profitable prey is significantly higher than that of the unprofitable prey — the predator will perceive both prey types as the more abundant profitable prey. However, when the two prey categories have a similar abundance, the predator will encounter an equal amount of both prey types; the resulting realized similarity will not be biased by prey abundance imbalance but instead be determined by the innate mimic–model morphological similarity. Finally, a third factor, *σ*_*j*_, controls the sensitivity of the predator recognition to the ratio of prey abundance. To represent this predator recognition process, we formulated the following abundance-dependent recognition function, which phenomenologically characterizes how the realized similarity is affected by the three aforementioned factors:

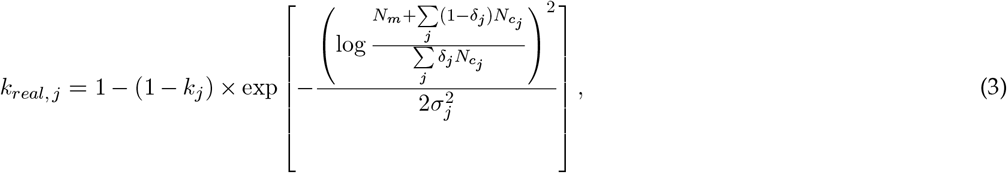

with *δ*_*j*_ = 0 if the profitability of 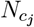 is equal or smaller than 0 and *δ*_*j*_ = 1 if the profitability is greater than 0; the term within the logarithm thereby represents the abundance ratio of the two prey categories. In a two-mimicry system that contains a Batesian mimic 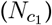 and a Müllerian mimic 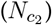, the abundance ratio term becomes 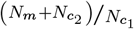. When the innate morphological similarity does not constrain realized similarity, i.e., easily distinguishable mimic with *k*_*j*_ = 0, *k*_*real, j*_ is solely determined by the abundance ratio of different prey categories and can vary freely between 0 and 1 (lower curve in Fig. 1C). As the innate morphological similarity *k*_*j*_ approaches 1 (i.e., mimics become hardly distinguishable), the realized similarity *k*_*rec, j*_ becomes less responsive to prey abundance ratio (upper curve in Fig. 1C). The functional form of eq. 3 implies that *k*_*real, j*_ approaches 1 when the abundance of the unprofitable and profitable prey is highly unbalanced (i.e., when 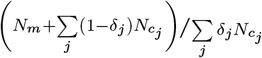 approaches 0 or *∞*). Under such a scenario, the predator treats the two prey categories as if both were the more abundant prey category: with optimal foraging, both are treated as the profitable prey if the abundance ratio approaches 0, and both are treated as the unprofitable prey if the ratio approaches *∞*.

#### Mimicry dynamic system

Finally, we incorporated optimal foraging theory, signal detection theory, and our novel abundance-dependent recognition function into a multi-mimicry dynamical system. We simulate the dynamics between mimics (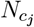 ; *j* = 1, 2, …, *n*), the model (*N*_*m*_), the predator (*N*_*p*_), and an alternative prey (*N*_*n*_):

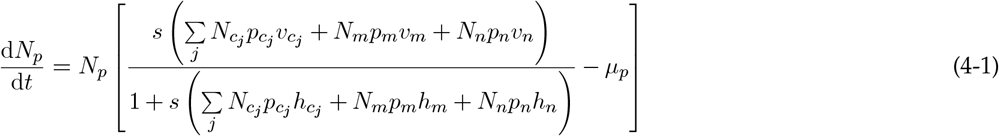

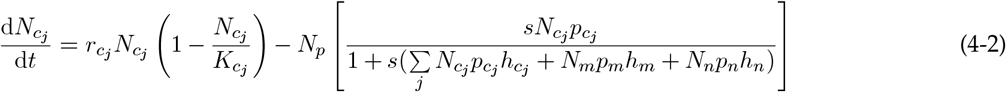

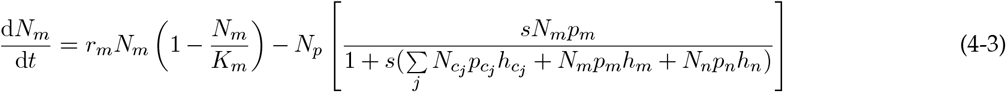

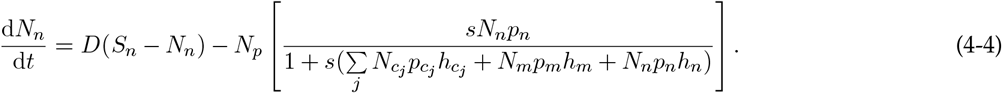

The per capita population growth rate of the predator consists of the consumption gain from optimal foraging (first term within the bracket in eq. 4-1) and the density-independent mortality rate *µ*_*p*_ (second term within the bracket in eq. 4-1). Specifically, the first term captures the total foraging gain the predator obtains from consuming various prey types, where the optimal combination of *p*_*i*_ is instantaneously adjusted at each time step to maximize the consumption gain (eq. 1) under the constraints imposed by signal detection theory (eq. 2) and abundance-dependent recognition (eq. 3). The mimics and the model grow logistically, with intrinsic growth rates 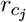 and *r*_*m*_, and carrying capacity 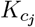 and *K*_*m*_, respectively. The population of the mimics and the model decreases due to predator consumption, which depends on the optimized probability 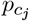 and *p*_*m*_, respectively (term within the bracket represents per predator consumption). Finally, we assumed that the alternative prey has an external population source, thus following chemostat dynamics with a continuous flux (*D*) and an external supply source (*S*_*n*_); the predator also attacks the alternative prey with optimized probability *p*_*n*_.

#### Numerical simulations

We simulated two different mimicry systems: (1) a single-mimic system with only the Batesian mimic (*N*_*c*_), and (2) a multi-mimic system with both Batesian mimic 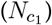 and Müllerian mimic 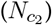. For simplicity, we omitted the secondary subscript *j* in 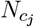 in the single-mimic model, but reintroduced it in the multi-mimic system (see Appendix A for the full equations). We used the fourth-order Runge–Kutta method from package deSolve (Karline Soetaert *et al*., 2010) to numerically solve our dynamical system. In all simulations, the integration step size was set to 0.02 and the dynamics were simulated for 12, 000 time steps. To incorporate optimal foraging, the values of *p*_*i*_ were adjusted at each integration step by finding the optimal combination that maximizes foraging gain (i.e., eq. 1). To better capture the nonlinear function form of eq. 2, the maximum foraging gain was found by evaluating eq. 1 with a range of *p*_*m*_ (and corresponding 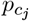) values from 0 to 1 in increments of 0.0001. While a larger increment for *p*_*m*_ (e.g., 0.1) could accelerate the numerical procedure, we note that it would lead to insufficient sampling along the nonlinear curve. Parameters in the single-mimic system are as follows (note again secondary subscripts *j* were omitted): *v*_*c*_ = 1.6, *v*_*n*_ = 0.8, *v*_*m*_ = 0, *h*_*c*_ = *h*_*n*_ = *h*_*m*_ = 1. The intrinsic growth rate and carrying capacity of the mimic and the model are *r*_*c*_ = *r*_*m*_ = 2 and *K*_*c*_ = *K*_*m*_ = 10, respectively, and the death rate of the predator is *µ*_*p*_ = 0.75. The influx of the alternative prey is controlled by parameters *S*_*n*_ = 20 and *D* = 1, and the search rate is *s* = 1. We used the following initial conditions for all simulations: *N*_*p*_(0) = 0.5, *N*_*c*_(0) = 1, *N*_*m*_(0) = 10, *N*_*n*_(0) = 20. The parameters for the multi-mimics system are set as follows (secondary subscripts *j* were reintroduced to distinguish the two mimics): 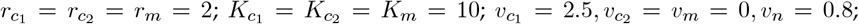 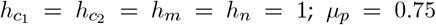, and *S*_*n*_ = 20. With 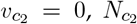 is the Müllerian mimic in the system, while 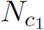 is the Batesian mimic with the highest profitability. The initial conditions for the multi-mimics system are identical to those of the single mimicry system, with 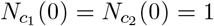.

To study the effect of mimic–model morphological similarity on community dynamics in the single-mimic system, we explored *k* across the parameter range of 0 to 1 by an interval of 0.01. For both the single-mimic and multi-mimicry systems, we studied the effect of abundance-dependent recognition (eq. 3) by varying the morphological similarity (i.e., *k* for the single-mimic system and both *k*_1_ and *k*_2_ for the multi-mimic system) from 0 to 1. All simulations were carried out using R 4.3.3 (R Core Team, 2024). When presenting simulation results, the last 80% of each simulated time series was used to calculate the mean and variance of each species’ abundance, which were then used to determine the stability and coexistence outcome of the system. Specifically, we set the extinction threshold for each species as 10^*−*10^. The stability threshold was set at 10^*−*12^, i.e., the system is considered stable if the variance of the predator population is less than 10^*−*11^. Since the predator interacts with all other prey and exhibits the highest population abundance, calculating the variance of the predator population is sufficient to determine the stability of the system.

## Result

### Single Batesian mimic system

#### Predator decision depending only on mimic–model morphological similarity

We first explored the dynamics of a single Batesian mimicry system where the predator’s decision depends only on the morphological similarity between the mimic and the model (*k*; Fig. 2A). We showed that along the spectrum of mimic–model similarity (*k* ranging from 0 to 1), low and high similarity produced cyclic dynamics while stable population dynamics were observed under intermediate similarity. Moreover, intermediate similarity (0.5 *< k <* 0.8) also resulted in higher predator and mimic abundance (red and orange, respectively, in Fig. 2A upper panel), suggesting that intermediate similarity benefits the mimicry system in terms of stability and species abundance. We discuss the dynamics below (see also Fig. 3 for detailed time series under different *k* values).

**Figure 2:**
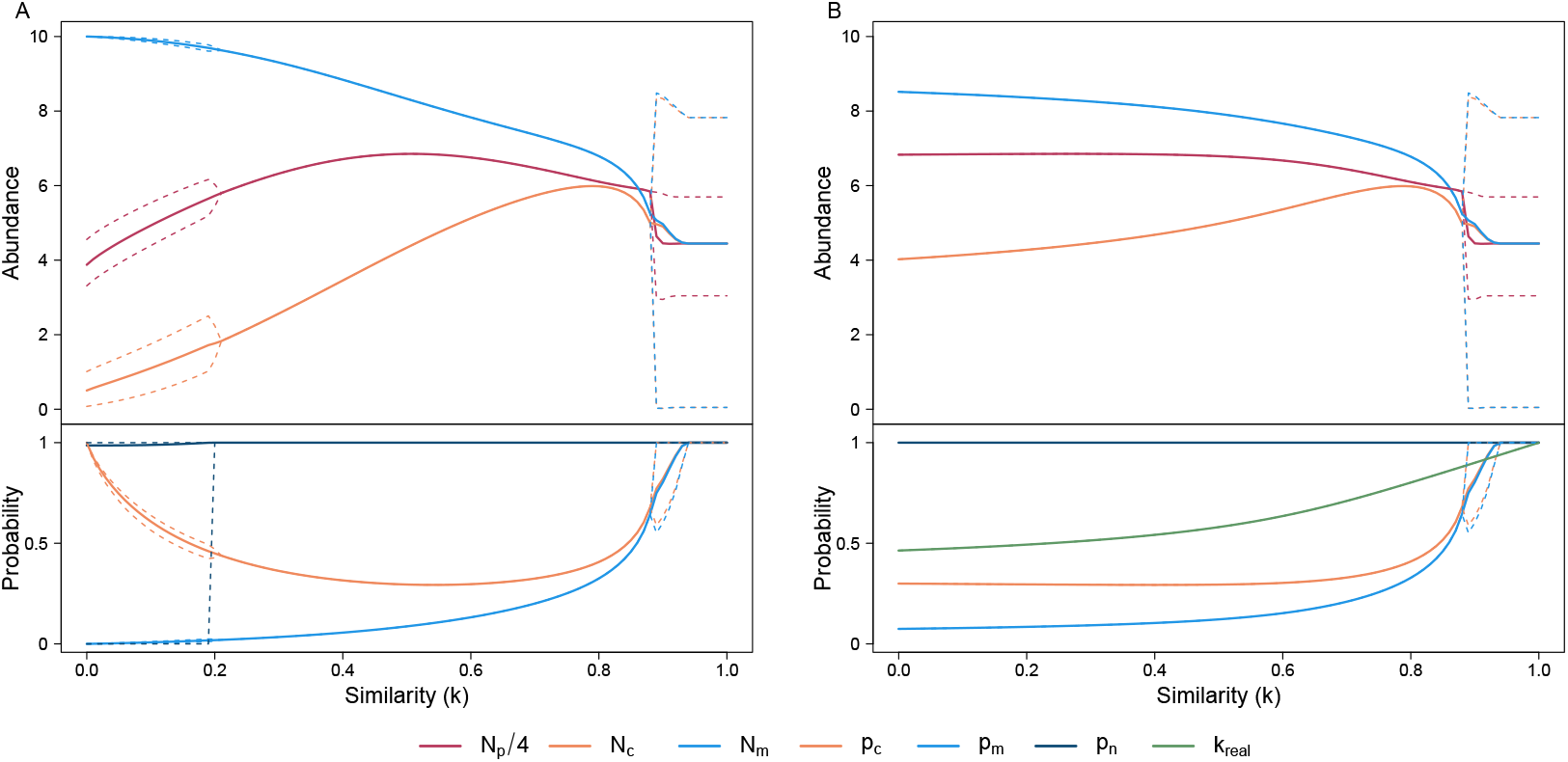
Bifurcation plot for the single mimic system, with predator recognition process based on (A) mimic–model morphological similarity or (B) abundance-dependent recognition. Both (A) and (B) bifurcate along the innate mimic–model morphological similarity (*k*). The upper panels of both plots illustrate the abundance of the predator (*N*_*p*_; red), the mimic (*N*_*c*_; orange), and the model (*N*_*m*_; light blue). Note that the abundance of the predator is divided by four for better visualization. The solid line represents the mean abundance, and the dashed line represents the maximum and minimum of the abundance. The lower panels of both plots illustrate the predator’s attack probability on the mimic (*p*_*c*_; orange), the model (*p*_*m*_; light blue), and the alternative prey (*p*_*n*_; dark blue). In panel (B), the realized recognition between the mimic and the model (*k*_*real*_; eq. 3) is plotted in green. The solid and dashed lines in the lower panels also represent the mean and maximum/minimum values, respectively.

**Figure 3:**
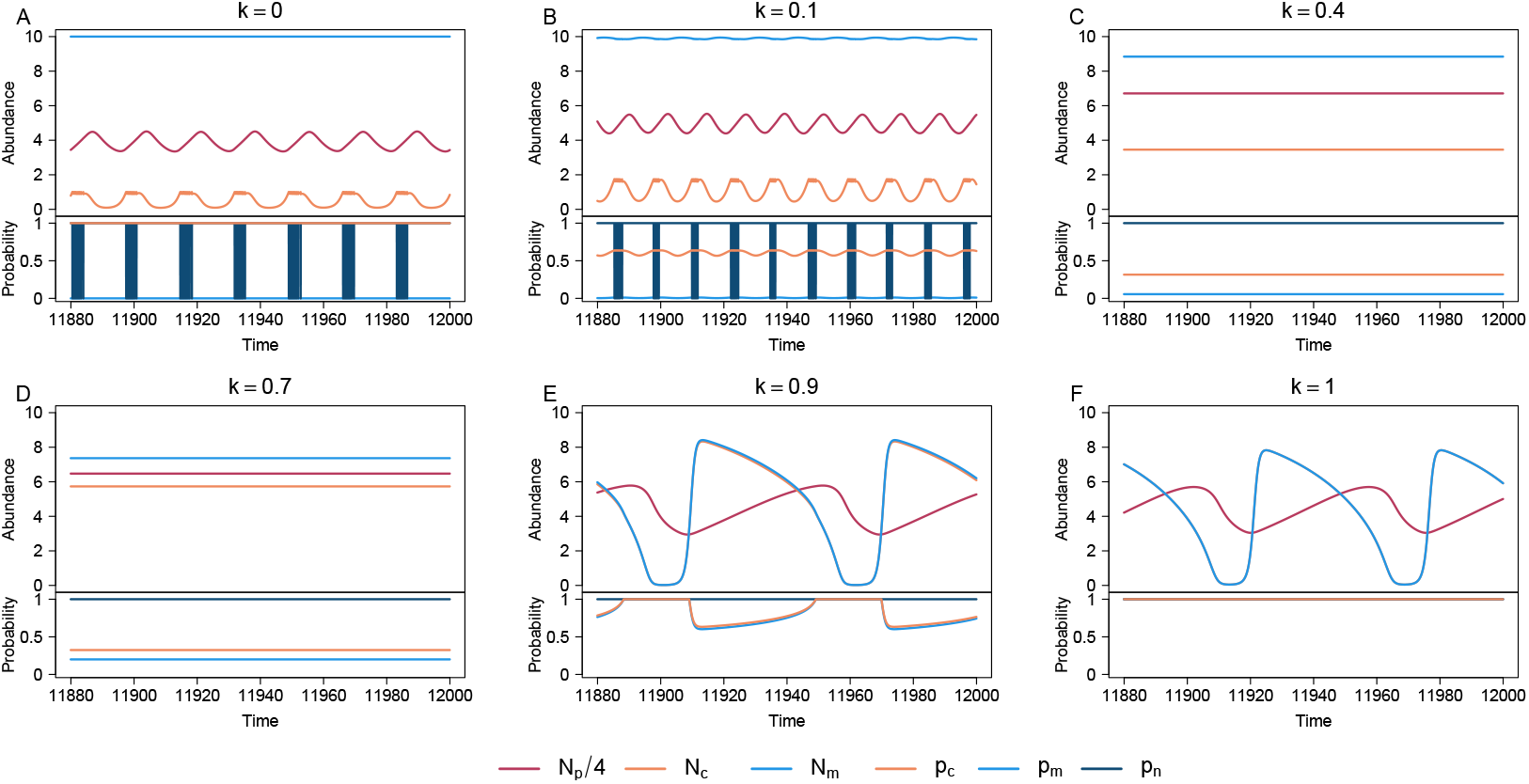
Time series of species abundance and predator attack probability for the single-mimic system with predator recognition based solely on mimic–model morphological similarity (*k*). Similarity (*k*) values vary as follows: (A) *k* = 0, (B) *k* = 0.1, (C) *k* = 0.4, (D) *k* = 0.7, (E) *k* = 0.9, and (F) *k* = 1. The upper half of each panel illustrates the abundance of the predator (*N*_*p*_; red), the mimic (*N*_*c*_; orange), and the model (*N*_*m*_; light blue). Note that the abundance of the predator is divided by four for better visualization. The lower half of each panel illustrates the predator’s probability of attacking the mimic (*p*_*c*1_; orange), the model (*p*_*m*_; light blue), and the alternative prey (*p*_*n*_; dark blue). Note in panel (F) the lines representing the mimic and the model completely overlap as they are perceived as the same species under *k* = 1.

When the similarity between the mimic and the model is zero (i.e., *k* = 0, left-most value in Fig. 2A), the mimic and the model are treated as two different species by the predator. In this scenario, cycles emerge from diet composition shifts: while predators always ignore the unprofitable model 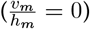 and always attack the mimic due to its high profitability, predators switch between including or excluding the alternative prey from its diet since changes in mimic abundance lead to different optimal *p*_*n*_ (Fig. 3A; see also Appendix B and Fig. S1 for analytical derivation). With slightly increased mimic–model similarity (i.e., increasing *k* towards approximately 0.2), the probability of the model being attacked increased while the attack probability of the mimic decreased (light blue and orange, respectively, in the lower panel of Fig. 2A). This is because predators now sometimes get confused between the mimic and the model and consume the wrong prey. Under such low similarity scenario (0 *< k <* 0.2), we observed that the system still exhibits cycles in its diet composition. However, unlike the scenario when *k* = 0, where predators exclude the model from their diet, here all three prey are included (Fig. 3B), demonstrating that even a slight similarity between the mimic and model can influence the community dynamics.

As the mimic–model similarity continues to increase (as *k* approaches 0.5), predators find it increasingly difficult to distinguish between the mimic and the model. Under this scenario, maintaining a high attack probability towards the mimic will lead to accidental consumption of the unprofitable model. To avoid this situation, optimal foraging leads predators to reduce their attack probability on the mimic. This foraging decision releases the predator from the cost of mistakenly consuming the model, ultimately stabilizing the predator population at higher abundance (Fig. 3C). At the same time, the mimic population increases in abundance due to the released predation pressure, while the model population continues to decline due to occasional predation.

Surprisingly, as the similarity between the mimic and the model increases further (0.6 *< k <* 0.8), the probability of attacking the mimic rises again, producing a U-shaped relationship between mimic–model similarity and attack probability across this intermediate range of *k*. This reversal occurs because, while predators could theoretically avoid accidentally consuming the model by further reducing attacks on mimics, optimal foraging prevents this strategy from being realized as the predator still requires foraging gain from the mimic–model species pair. Instead, predators unavoidably need to increase their attack probability on both prey items to make up for the frequent unprofitable accidental attacks, thereby decreasing the abundance of the predator. Initially, this decline in predator abundance allows the mimic population to rise further, reaching its peak just before similarity exceeds *k* = 0.8 (Fig. 3D). However, once similarity surpasses this threshold, the rising attack probability on mimics leads to a decrease in mimic abundance. Consequently, the peaks in predator and mimic abundance occur at different values of *k*, reflecting the shifting balance between prey profitability and foraging pressure within the mimicry complex.

In addition to the counterintuitive increase in *p*_*c*_, a higher mimic–model similarity also results in the reoccurrence of cyclic dynamics (0.88 *< k <* 1; right-most region in Fig. 2A). When 0.88 *< k <* 0.94, population cycles are characterized by diet composition shifts (Fig. 3E) similar to those seen under low mimic–model similarity. However, once similarities surpass a certain threshold (*k >* 0.93), the cycles are akin to classic predator–prey population cycles and no longer involve diet composition shifts. Instead, predators now view the mimic–model species pair as a single species and consistently include both in their diet (*p*_*m*_ = *p*_*c*_ = 1; Fig. 3F). Note that despite their inability to distinguish the mimic and the model apart, predators do not discard the mimic–model species pair from their diet because its collective value remains profitable. We show in Figure S2 that making *v*_*m*_ *<* 0 can, heuristically, reduce the collective profitability of the mimic–model species pair, eventually causing predators to discard the species pair altogether if they share high morphological similarity (Fig. S2).

#### Predator decision depending on abundance-dependent recognition

We next relaxed the assumption that the predator’s decision process is solely determined by mimic– model morphological similarity (*k*), making it dependent on the abundance ratio of two prey categories, i.e., the profitable mimic and the unprofitable model (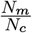 ; eq. 3). We used the notation *k*_*real*_ to represent the realized similarity between the mimic and the model from the perspective of the predator (again, secondary subscript omitted for simplicity). Unlike the previous scenario (Fig. 2A), which produced diet shift cycles under low similarity values, abundance-dependent recognition leads to stable dynamics under a wide range of innate similarity values (0 *< k <* 0.88; Fig. 2B). This suggests that when the innate mimic–model similarity is not too high, the predator recognition is influenced by the observed 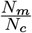 ratio and reaches an intermediate realized similarity (green line in Fig. 2B lower panel). The realized intermediate similarity (0.46 *< k*_*real*_ *<* 0.88) corresponds to values in Figure 2A that stabilize the system and lead to higher predator and mimic abundance. In other words, abundance-dependent recognition leads predators to reduce attack on mimics with low morphological similarity, causing the system to exhibit stable dynamics associated with intermediate similarity values (i.e., *k*_*real*_ *> k*).

However, the predator becomes less responsive to the prey abundance ratio, and the realized similarity becomes increasingly similar to *k*, as predator recognition becomes increasingly constrained by the innate mimic–model similarity (i.e., high *k*). That is, the predator cannot adopt its realized similarity to an intermediate value that would stabilize the system. Instead, the high innate mimic–model morphological similarity causes the predator to perceive the two prey as a single species, which causes 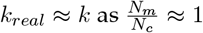; the resulting dynamics is a cyclic behavior with reduced population sizes (corresponding to a high *k* scenario in Fig. 2A).

### Multi-mimic system with abundance-dependent recognition

We expanded our model to consider a multi-mimic mimicry system consisting of a Batesian mimic 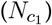, a Müllerian mimic 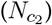, and a model species (*N*_*m*_), with the latter two considered as unprofitable prey 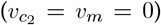. We considered this community composition as it represents the simplest mimicry complex with multiple types of mimicries (see also Fig. S3 for an example with more species). We explored how the Batesian mimic–model morphological similarity (*k*_1_) and the Müllerian mimic–model morphological similarity (*k*_2_) influenced community dynamics, assuming the predator possesses abundance-dependent recognition (see also Fig. **??** for the case where foraging decisions depend solely on mimic–model morphological similarity). Intuitively, a high *k*_*real*, 1_ value causes the predator to perceive the mimicry complex as a more profitable group of prey, whereas a high *k*_*real*, 2_ would decrease the overall profitability of the mimicry complex. Briefly, the simulation reveals that while intermediate similarity in both Batesian and Müllerian mimicry stabilizes the system and increases the abundance of the predator and mimics, different types of mimicry vary in their impact on the species abundance (Fig. 4).

**Figure 4:**
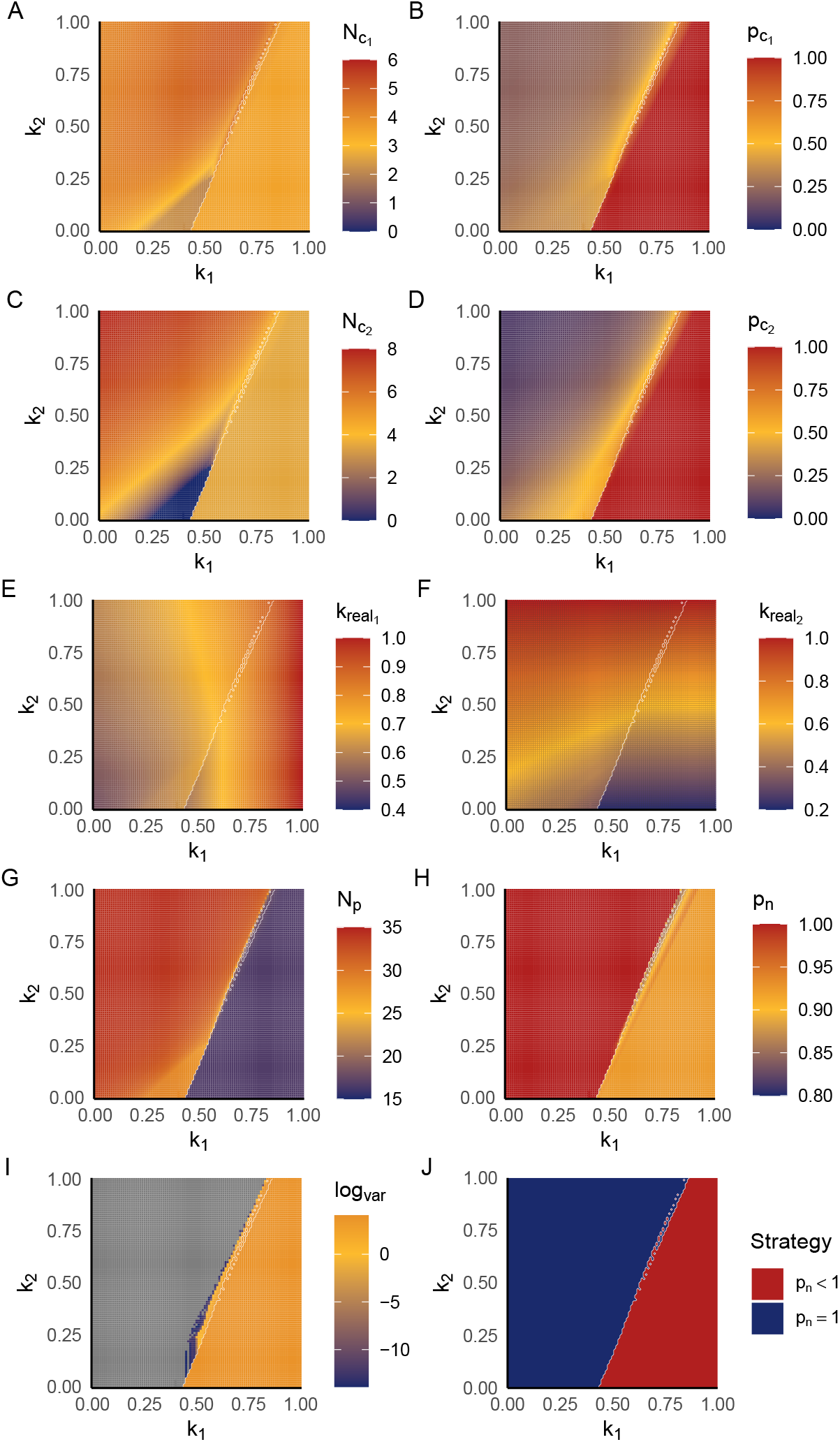
The effect of the Batesian mimic–model morphological similarity (*k*_1_; x-axis) and the Müllerian mimic–model morphological similarity (*k*_2_; y-axis) on system dynamics and species abundances. The different panels represent different variables: (A) Batesian mimic abundance 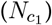, (B) attack probability on the Batesian mimic 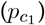, (C) the Müllerian mimic abundance 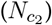, (D) attack probability on the Müllerian mimic 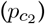, (E) realized Batesian mimic similarity (*k*_*real*, 1_), (F) realized Müllerian mimic similarity (*k*_*real*, 2_), (G) predator abundance (*N*_*p*_), (H) attack probability on the alternative prey (*p*_*n*_), and (I) the log(variance) of the predator population fluctuation, serving as an indicator of community stability. Panel (J) depicts whether *p*_*n*_ = 1 (blue) or *p*_*n*_ *<* 1 (red), which serves as a binary indicator of the two dynamical regimes: (1) the consistent alternative prey inclusion regime (blue) and (2) the occasional alternative prey exclusion regime (red). The white outline in all panels depicts the boundary separating the two dynamical regimes in (J). Note that the abundances of different state variables are on different scales to better show the pattern of each variable.

Our simulation suggests that the system exhibits two distinct dynamical regimes, governed by the interplay between *k*_1_ and *k*_2_: (1) a consistent alternative prey inclusion regime (top-left of Fig. 4) and (2) an occasional alternative prey exclusion regime (bottom-right of Fig. 4). In the first regime, in addition to always consuming the alternative prey (average *p*_*n*_ = 1; Fig. 4H, J), it is characterized by lower 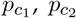 (top-left of Fig. 4B, D) and higher abundance of 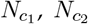, and *N*_*p*_ (top-left of Fig. 4A, C, E). In contrast, in the occasional alternative prey exclusion regime, the alternative prey is not always included (average *p*_*n*_ *<* 1; Fig. 4H, J) but the Batesian and the Müllerian mimics are consistently included in the diet with high attack probability, leading to their lower abundance (lower-right of Fig. 4A–D). The difference in the diet composition suggests that these two regimes are governed by the predator’s diet strategy (see Appendix B for the analytical derivation of this boundary).

Additionally, most scenarios in the top-left consistent alternative prey inclusion regime exhibit stable population dynamics, while all scenarios in the lower-right occasional alternative prey exclusion regime exhibit cyclic dynamics (Fig. 4I). The boundary separating the two regimes shows a positive association between *k*_1_ and *k*_2_, indicating that a higher value of *k*_2_ allows the system to tolerate a greater *k*_1_ before becoming destabilized (Fig. 4I). When *k*_1_ is sufficiently high, the system eventually becomes unstable as the predator increasingly perceives the mimicry complex as a profitable food source, akin to the instability seen in the single Batesian mimic system with high similarity (Fig.2B). However, the presence of a Müllerian mimic can mitigate this effect: a Müllerian mimic with sufficiently high *k*_2_ can stabilize the system, especially when *k*_1_ is fixed at low to intermediate values (Fig. 4I). This demonstrates that mimic types differ in their impact on system stability, with a perfect Batesian mimic (high *k*_1_) causing the system to be unstable and a perfect Müllerian mimic (high *k*_2_) promoting stability.

Beyond stability, we examined how the Batesian mimic–model morphological similarity (*k*_1_) affects mimic abundance, while keeping the Müllerian mimic’s morphological similarity (*k*_2_) constant. Under abundance-dependent recognition, *k*_*real*, 1_ remains within an intermediate range (0.5 *< k*_*real*, 1_ *<* 0.8; Fig. 4E), and Batesian mimic abundance exhibits a hump-shape pattern with increasing *k*_1_ (orange line in Fig. 5A). Moreover, compared to the scenario where predator recognition relies only on morphological similarity (Fig. **??**A), the resulting intermediate *k*_*real*, 1_ leads to a higher Batesian mimic abundance. The Müllerian mimic abundance, on the other hand, decreases monotonically with the increasing *k*_1_ (green line in Fig. 5A). As *k*_1_ further increases, the dynamics eventually transition into the aforementioned occasional alternative prey exclusion regime, characterized by unstable dynamics with consistently low abundance of both mimics as the predator now consumes the entire mimicry complex as a single palatable prey.

**Figure 5:**
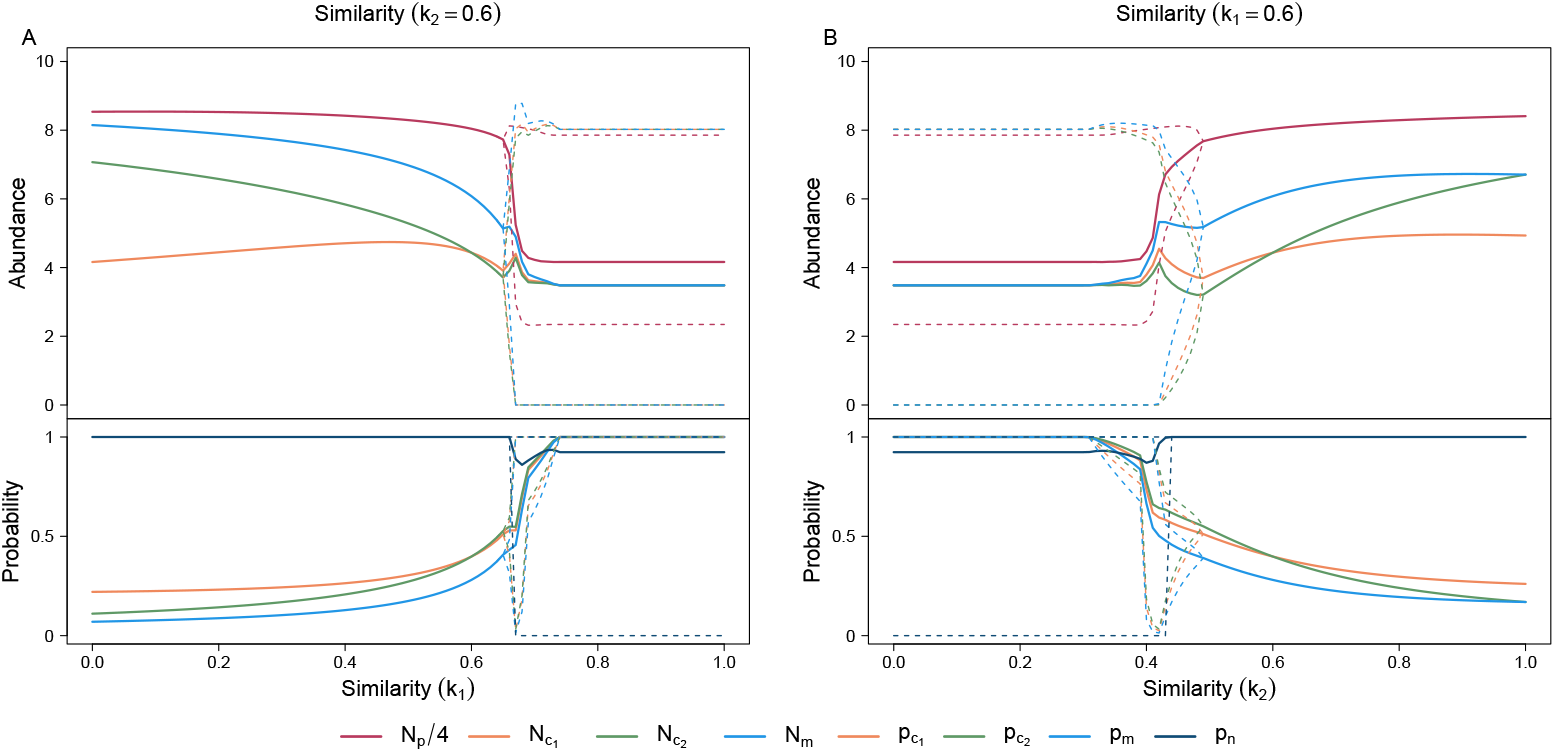
Bifurcation plot of the multi-mimicry system. (A) bifurcation along the Batesian mimic–model morphological similarity (*k*_1_) while fixing the Müllerian mimic’s morphological similarity (*k*_2_ = 0.6). (B) bifurcation along the *k*_2_ while fixing *k*_1_ = 0.6. The y-axis of the upper panels shows abundance, and the lower panels show the attack probability. The upper panels of both plots illustrate the abundances of the predator (*N*_*p*_; red), the Batesian mimic (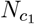; orange), the Müllerian mimic (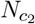 ; green), and the model (*N*_*m*_; light blue). Note that the abundance of predator is divided by four for better visualization. The lower panels of both plots illustrate the predator’s attack probability on the Batesian mimic (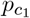 ; orange), the Müllerian mimic (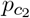 ; green), the model (*p*_*m*_; light blue), and the alternative prey (*p*_*n*_; dark blue). The solid and dashed lines in all panels represent the mean and maximum/minimum values, respectively.

Finally, we examined how varying the Müllerian mimic–model morphological similarity (*k*_2_) affects mimic abundance, while keeping the Batesian mimic’s morphological similarity (*k*_1_) fixed. When the Batesian mimic has an innate intermediate similarity (0.5 *< k*_1_ *<* 0.8), the system is likely within the occasional alternative prey exclusion regime with low abundance of both mimics (Fig. 4A, C). Under this scenario, a Müllerian mimic with high similarity provides protection to the mimicry complex: higher *k*_2_ balances out the impact from a highly similar Batesian mimic, resulting in the transition into the other dynamical regime with high mimic abundances. Moreover, within the consistent alternative prey inclusion regime, while *k*_*real*, 1_ settles within an intermediate level (0.5 *< k*_*real*, 1_ *<* 0.8) and the Batesian mimic thereby maintains higher abundance, *k*_*real*, 2_ increases monotonically with increasing *k*_2_ (Fig. 4F). As a result, Müllerian mimic abundance increases with *k*_2_ (Fig. 5B), suggesting that the Müllerian mimicry provides more protection under higher morphological similarity. Together, these results underscore the contrasting ecological roles of the two mimic types. For the Batesian mimic, intermediate similarity to the model reduces predation while avoiding the high *k*_1_ that causes predators to consume the entire mimicry complex as a single profitable prey. In contrast, Müllerian mimics benefit from high similarity as it reinforces predator learning, thereby leading to a steady increase in abundance with increasing *k*_2_.

## Discussion

We showed that imperfect mimicry (i.e., intermediate similarity between the mimic and the model) promotes stable ecological dynamics and allows the mimic to reach high abundance in a single Batesian mimicry system. This outcome holds whether predator recognition is shaped solely by morphological similarity or by a combination of morphological similarity and prey abundance ratio (Fig. 2). In a multi-mimicry system including both Batesian and Müllerian mimics, intermediate similarity remains beneficial for the Batesian mimic, while high similarity for the Müllerian mimic leads to increased mimic abundance (Fig. 4). These contrasting patterns suggest that the two mimic types have different impacts on the system: the Batesian mimic can destabilize the system (and lead to low mimic abundance) when too similar to the model, whereas the Müllerian mimic with high similarity can promote system stability (and lead to high mimic abundance). Together, our results suggest that imperfect mimicry can lead to ecologically stable dynamics across different mimicry systems, offering new insights into how mimicry persists in nature.

Our results provide a mechanistic perspective on the “eye of the beholder hypothesis” (Cuthill & Bennett, 1993, Dittrich *et al*., 1993), which proposes that mimicry deemed imperfect by human observers may be functionally effective from the perspective of the predator. Specifically, our abundance-dependent recognition framework demonstrates that predators tend to perceive a higher level of realized similarity than what is suggested by morphology alone (i.e., *k*_*real, j*_ *≥ k*_*j*_). Our formulation of abundance-dependent recognition echoes Müller’s original idea that the inclusion of more unprofitable prey strengthens the protective benefits of aposematic signals (Müller, 1879), and it aligns with the principles of Pavlovian conditioning, in which repeated signal exposures reinforce learned associations. Our abundance-dependent recognition also accommodates predator sensitivity to various prey categories through the parameter *σ*_*j*_ (Fig. S5). By incorporating prey abundance into predator recognition and mimicry dynamics, our framework extends beyond static morphological similarity and highlights an ecological dimension that human observers typically overlook. This mismatch between predator and human perception helps explain how the “eye of the beholder” effect could emerge from predator decision–making behavior.

Our results from the single Batesian mimicry system illustrates that imperfect mimicry could promote mimic abundance, leading to the prediction that mimics in simple Batesian systems should generally exhibit intermediate resemblance to their model. This aligns with recent empirical findings showing that imperfect mimicry is more evolutionarily stable than perfect mimicry (Kelly *et al*., 2025). In contrast, our results from the multi-mimicry system suggest that while Müllerian mimics alone should achieve higher similarity, the interplay between different mimic types can generate more complicated dynamics. Empirical studies also indirectly support this idea: Müllerian mimicry rings could include dozens of species with varying degrees of resemblance and even overlapping traits across different mimicry rings (Wilson *et al*., 2012, Wilson *et al*., 2022, Motyka *et al*., 2020), potentially due to the existence of Batesian mimics or quasi-Müllerian mimics in the mimicry complex. Therefore, based on our theoretical results, we hypothesize that Müllerian mimics in nature may exhibit a greater diversity of aposematic signals when Batesian mimics or quasi-Müllerian mimics are also involved. This prediction can be tested empirically by comparing the trait diversity among Müllerian mimics across different multi-mimicry complexes.

Our predictions regarding the benefits of imperfect mimicry can be tested by meta-analysis of behavior experiments and comparative analysis of trait data. For example, a meta-analysis on mimic–model morphological similarity could examine if Müllerian mimics typically exhibit a higher similarity with the model, while single Batesian mimicry systems are more likely to exhibit intermediate similarity with the model. One can analyze morphological traits such as color, pattern, or other aposematic signals to characterize the similarity between mimics and models (Eliason *et al*., 2019, Maia *et al*., 2019, Kelly *et al*., 2021), thereby providing a basis to evaluate whether empirical systems align with theoretical predictions. In addition, combining trait data with phylogenetic information can reveal whether the aposematic signal is under selection (Eliason *et al*., 2019). In Müllerian mimicry systems, we expect strong directional selection on these traits with model and mimic converging to one aposematic signal. In contrast, we expect predation-driven selection in Batesian mimicry systems to stabilize traits at intermediate similarity. These predictions can be tested through behavior experiments, such as assessing predator attack probabilities on an array of mimics with various degrees of similarity (Tseng *et al*., 2014); our predictions will be supported if the mimic with intermediate similarity suffers the least attack.

Finally, we encourage future work to explicitly integrate aposematic traits into the theoretical framework of mimicry dynamics (Holen & Johnstone, 2004, Tomizuka & Tachiki, 2024). One promising approach is to replace the similarity parameter in our model with species trait distance in a multi-dimensional trait space. This trait-explicit framework would provide a more mechanistic interpretation of the mimic–model similarity parameter and avoid potential artifacts arising from the phenomenological linkage between attack probabilities in signal detection theory. For instance, in our current framework, the attack probabilities of the two mimics are indirectly constrained by their linkage to the model’s attack probability (see Appendix C), which can create abrupt abundance drops at low similarity values that do not reflect biologically optimal predator behavior (Fig. 4C). Furthermore, a trait-explicit framework allows the investigation of eco-evolutionary feedback dynamics (Tomizuka & Tachiki, 2024), where mimic trait distributions evolve under selection imposed by predators. Such a framework could further help determine not only the ecological stability but also the evolutionary persistence of imperfect mimicry. Overall, our study underscores the significance of considering predator recognition and population dynamics into mimicry theory, offering new insights into how imperfect mimicry can be maintained in natural, complex mimicry systems.

## Acknowledgements

We thank Chun-Wei Chang, Hsi-Cheng Ho, Chi-Yun Kuo, Hui-Yun Tseng, Ching-Lin Huang, Yu-Pei Tseng, Joe Wan, Shan-Min Chen, Yu-Chen Chen, and members of the Ke Lab for comments on early drafts of the manuscript. This study is supported by the Taiwan Ministry of Education Yushan Fellow Program (MOE-110-YSFAG-0003-001-P1) and the Taiwan National Science and Technology Council (MOST 111-2621-B-002-001-MY3 and NSTC 114-2628-B- 002-023-).

## Supplementary information for

## Appendix A Equations for the single and the two-mimic system

The single-mimic system contains a Batesian mimic (*N*_*c*_), model (*N*_*m*_), predator (*N*_*p*_), and the alternative prey (*N*_*n*_). The explicit equations for the single-mimic system are as follows (note that the secondary subscript *j* in 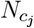 and associated parameters are omitted for simplicity):

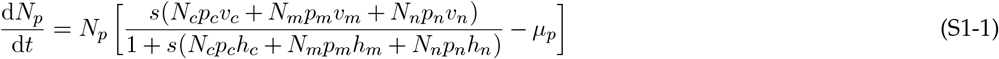

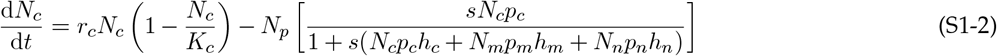

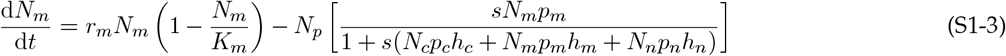

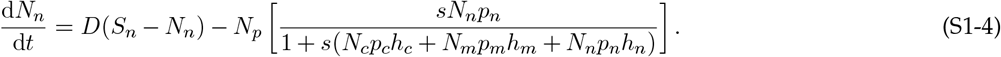

The two-mimic system contains two types of mimics (secondary subscript *j* reintroduced to distinguish them), with 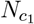 as the Batesian mimic and 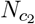 as the Müllerian mimic. The explicit equations for the two-mimic system are as follows:

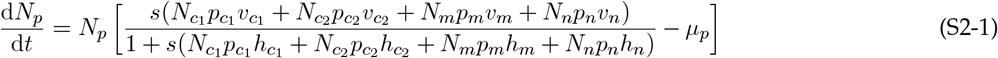

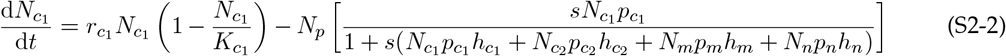

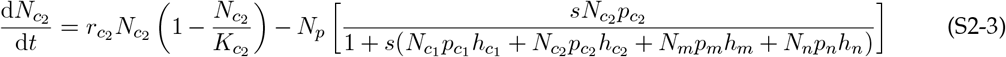

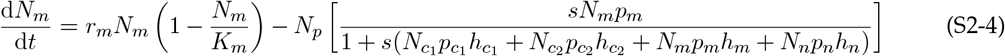

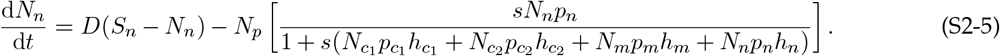

## Appendix B Analytical criteria for diet shifting

Optimal foraging theory is a key component of our theoretical framework, producing cycling behavior in both single Batesian mimicry and multi-mimic systems. Here we present the analytical criteria and simulation results for this cycling behavior in different systems.

### Diet shift criterion for single Batesian mimic system

In the single Batesian mimicry framework, we illustrated two mechanisms that produce cycles: diet shift and predator–prey interaction. In Fig. 2A, we show that a diet shift cycle could happen when *k <* 0.2 and 0.88 *< k <* 0.94. Here, for the single Batesian mimicry system, we derive the analytical criterion for predators to include the alternative prey in their diet when the mimicry complex is determined to be the more profitable food. The energy gained by predators under different diet compositions is as follows:

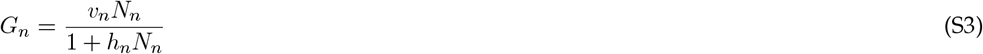

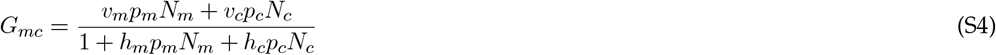

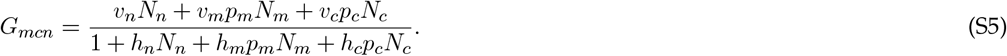

Here, *G*_*n*_, *G*_*mc*_, and *G*_*mcn*_ represent the energy gain when the predator diet consists of only the alternative prey, only the mimicry complex, and all three prey items, respectively. For predators to include alternative prey into the diet when it currently only consumes the mimicry complex, the gain from including all three species has to be greater than that when only including mimics and models, i.e., *G*_*mcn*_ *> G*_*mc*_. By rearranging the terms, we arrive at the following criterion:

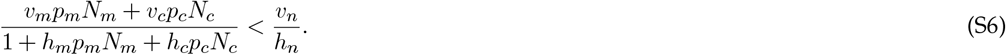

This result shows that switching from *G*_*mc*_ to *G*_*mcn*_ requires the profitability of the alternative prey to be greater than the energy gain obtained from only consuming the mimicry species pair. In Figure S1, we show a time series illustrating the diet shift from *G*_*mcn*_ to *G*_*mc*_ (i.e., *p*_*n*_ switches between 1 to 0, respectively) when *k* = 0.1 (see also Fig. 2A). Notably, we depict our analytical derivation with gray strips in Figure S1, showing that the analytic criterion matches the time points of diet switching in the simulation result.

Besides showing the criterion for a predator to switch from only consuming the mimicry complex to consuming all three prey species (i.e., the inclusion of the alternative prey), we also provide the criterion for when a predator will switch its diet from consuming only alternative prey to consuming all three prey species (i.e., the inclusion of the mimicry pair). The criterion for the predator to switch diet is *G*_*mcn*_ *> G*_*n*_, which corresponds to:

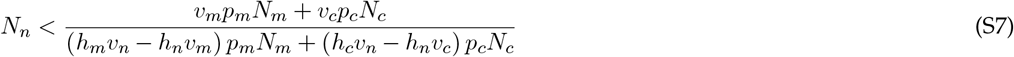

### Regime shift criterion for multi-mimic system

For the multi-mimic system, Figure 4 shows that there is a boundary separating two dynamical regimes. The two regimes are characterized by whether the alternative prey is always included in the predator’s diet or not (Fig. 4J). In the consistent alternative prey inclusion regime (top-left of Fig. 4), the energy gain can is as follows:

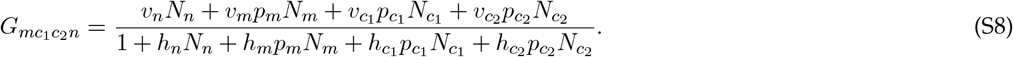

We could also write out the energy gain term in the occasional alternative prey exclusion regime (bottom-right of Fig. 4), where the predator always incorporates the mimicry complex but does not always incorporate the alternative prey:

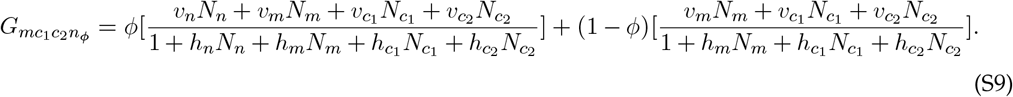

Here, *ϕ* is the proportion of time that the alternative prey is included in the diet, which can be obtained by the long-term average of *p*_*n*_ since it is a binary variable. When 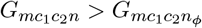, the system enters the consistent alternative prey inclusion regime with stable dynamics. Conversely, when 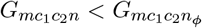, the system enters the occasional alternative prey exclusion regime with unstable dynamics. Note that the white outline in Fig. 4 was drawn based on the above criterion, with *ϕ* equal to the average *p*_*n*_ obtained from the simulation.

## Appendix C Caution against multi-species signal detection theory

In the multi-mimicry system, we incorporated signal detection theory for the two mimic species independently as:

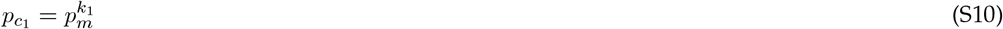

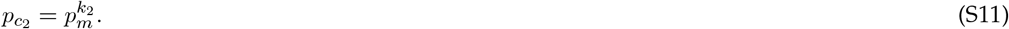

This expression could be rewritten as:

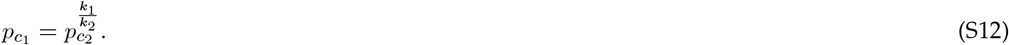

Under this expression, numeric artifacts may occur when both *k*_1_ and *k*_2_ are low as the attack probabilities of the two mimics are indirectly constrained. Consider the case where *k*_1_ = 0 and *k*_2_ = 0, which represents the scenario in which the predator could perfectly distinguish between the Batesian mimic, the Müllerian mimic, and the model. When the predator is capable of distinguishing between all prey items, one would intuitively expect that predators should only attack the Batesian mimic; indeed, the optimal foraging theory results in *p*_*c*_1 = 1. However, with *p*_*m*_ becoming 0 given *k*_1_ = 0, the fact that *k*_2_ = 0 numerically guarantees that 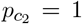 (lower-left region of Figs 4C and **??**C). This conflict between biological intuition and numeric results comes from the direct multi-species extension of the signal detection theory, and can be seen in the lower left of Figs 4 and **??**. We note that abundance-dependent recognition can partially resolve this numerical artifact, since the realized similarity *k*_*real, j*_ will deviate from the low innate morphological similarity and achieve an intermediate value. To further resolve this artifact, a trait-based approach could be a promising alternative approach to model multi-mimicry systems, as mentioned in the Discussion.

## Supplementary Figures

**Figure S1:**
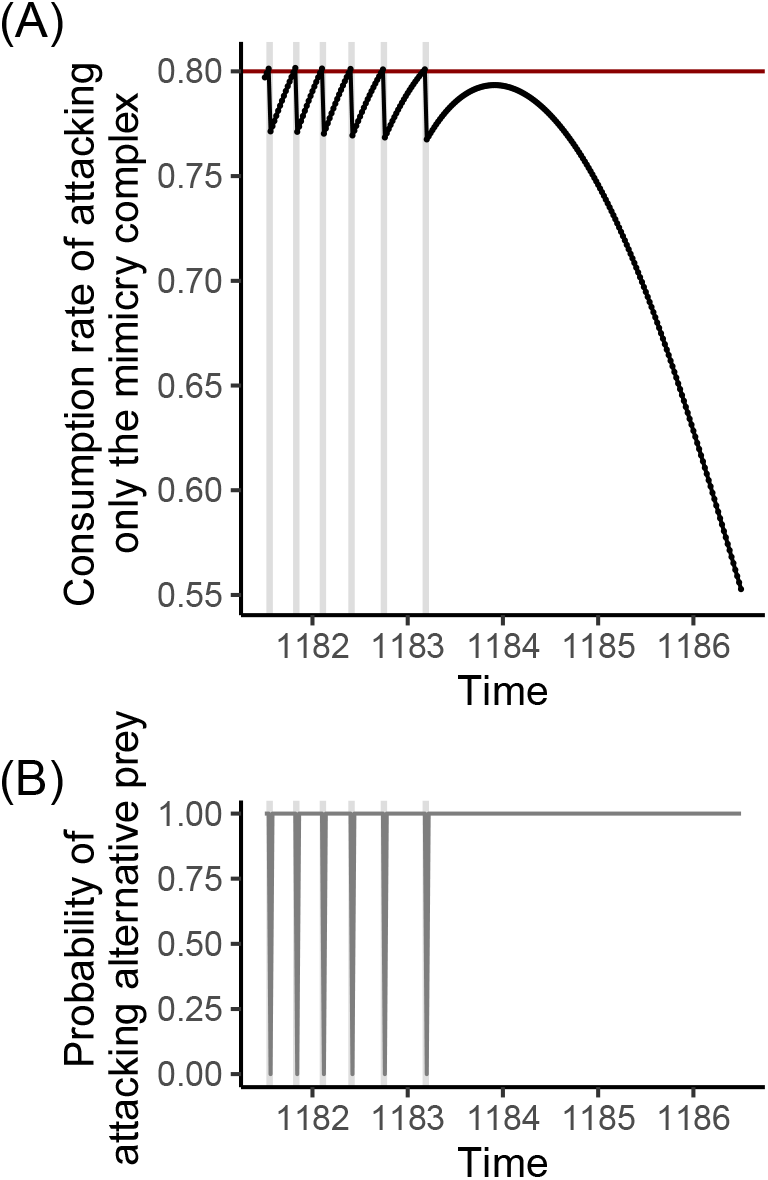
Time series plots showing the (A) diet switching criterion and (B) probability of consuming the alternative prey (*p*_*n*_) in a single Batesian mimicry system with *k* = 0.1. Here, we show the diet switching behavior from only consuming the mimicry species pair (*G*_*mc*_) to including the alternative prey (*G*_*mcn*_). The x-axis shows time step 1181 to 1187 (with integration step size 0.02). The black line in panel A is the consumption rate of attacking only the mimicry complex and the red line shows the profitability of the alternative prey (constant 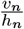). The blue line in panel B is the probability of attacking the alternative prey (*p*_*n*_). Grey strips indicate the time period when the switching criterion is met, thereby inducing a diet switch. The parameter values are identical to those in the main text, see section Numerical simulations for details.

**Figure S2:**
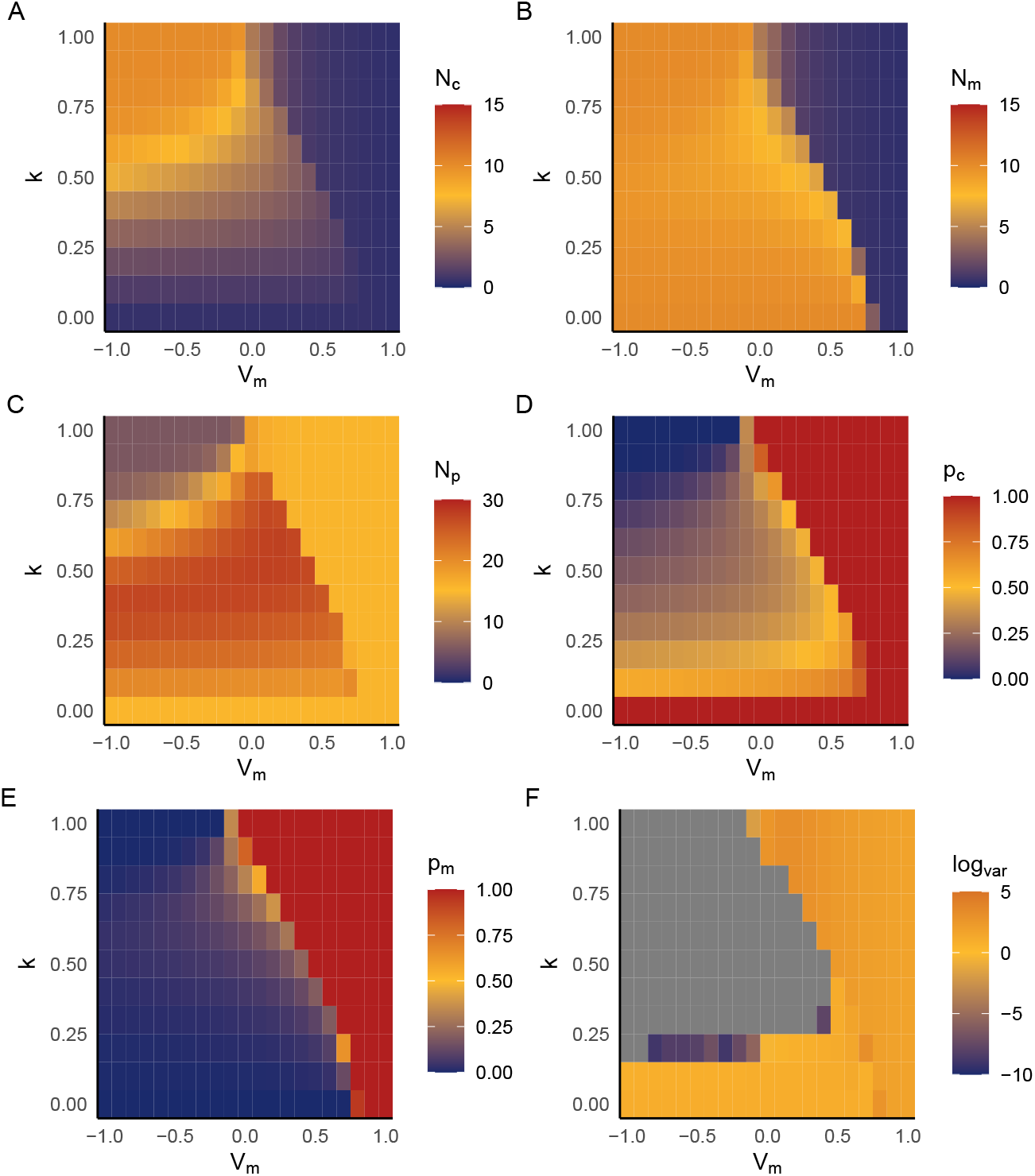
The effect of the model species value (*v*_*m*_; x-axis) and the mimic–model similarity (*k*; y-axis) on the single Batesian mimicry system without abundance-dependent similarity. A lower model value indicates a lower profitability of consuming the model, which also influences the collective profitability of the mimic–model species pair; note the default value in the main text is *v*_*m*_ = 0. Different panels represent different variables: (A) mimic abundance (*N*_*c*_), (B) model abundance (*N*_*m*_), (C) predator abundance (*N*_*p*_), (D) the attacked probability on the mimic (*p*_*c*_), (E) the attacked probability on the model (*p*_*m*_), and (F) the log(variance) of the predator abundance fluctuation, serving as an indicator of whether the system is cycling or not. See section Numerical simulations for parameter details.

**Figure S3:**
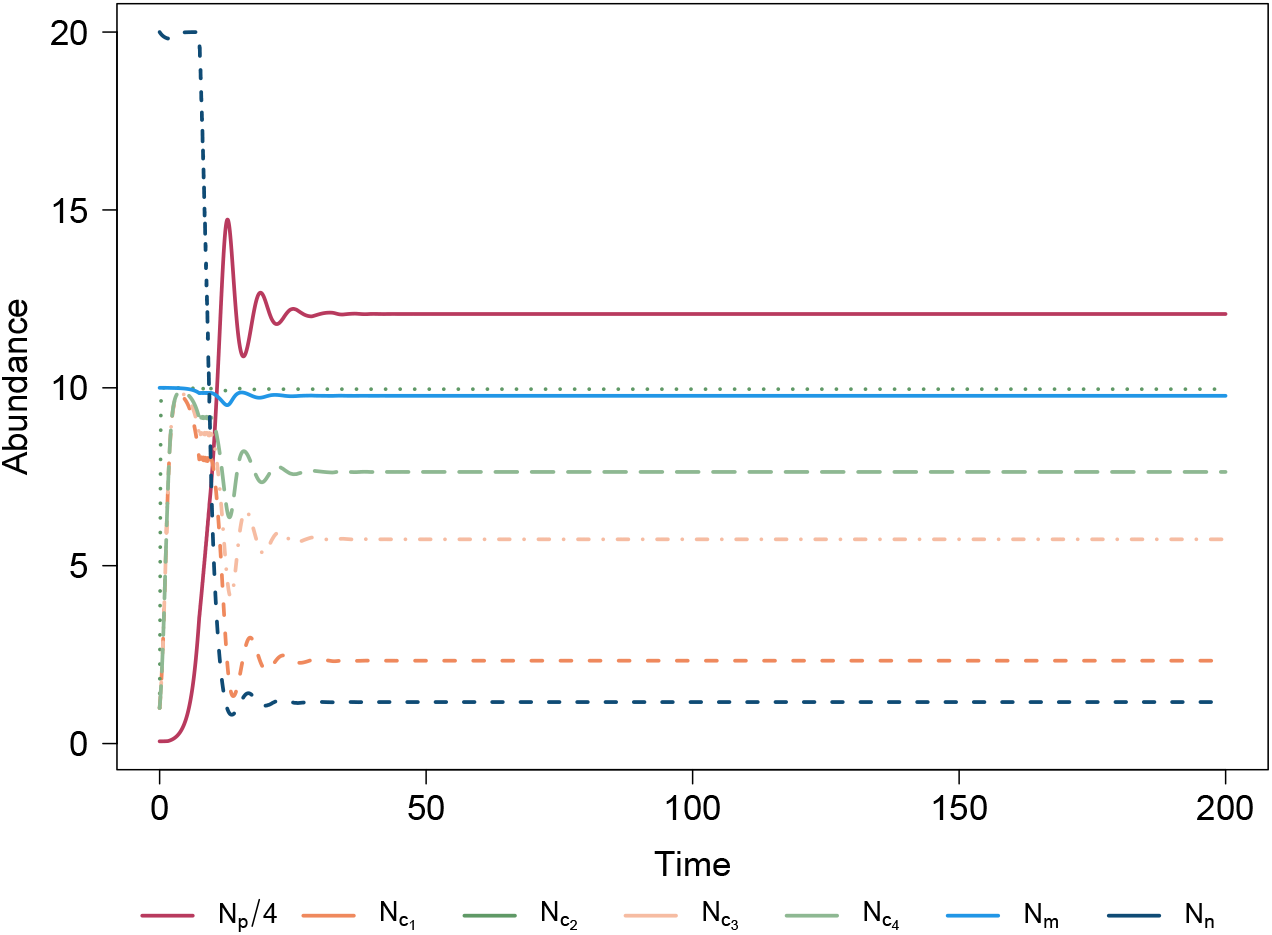
Time series of species abundances in a four-mimicry system where predator recognition is based solely on mimic–model morphological similarity 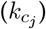. The system includes: two Batesian mimics (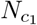 and 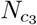 ; orange and light orange, respectively) and two Müllerian mimics (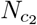 and 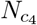 ; green and light green, respectively), the model (*N*_*m*_; light blue), the predator (*N*_*p*_; red), and the alternative prey (*N*_*n*_; dark blue). Note that the abundance of predators is divided by four for better visualization. Similarity 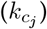 values as follows: 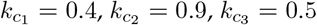, and 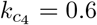. Other parameter follow the main text, with additional parameters for 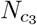 and 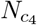 including: 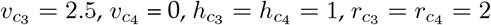, and 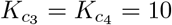.

**Figure S4:**
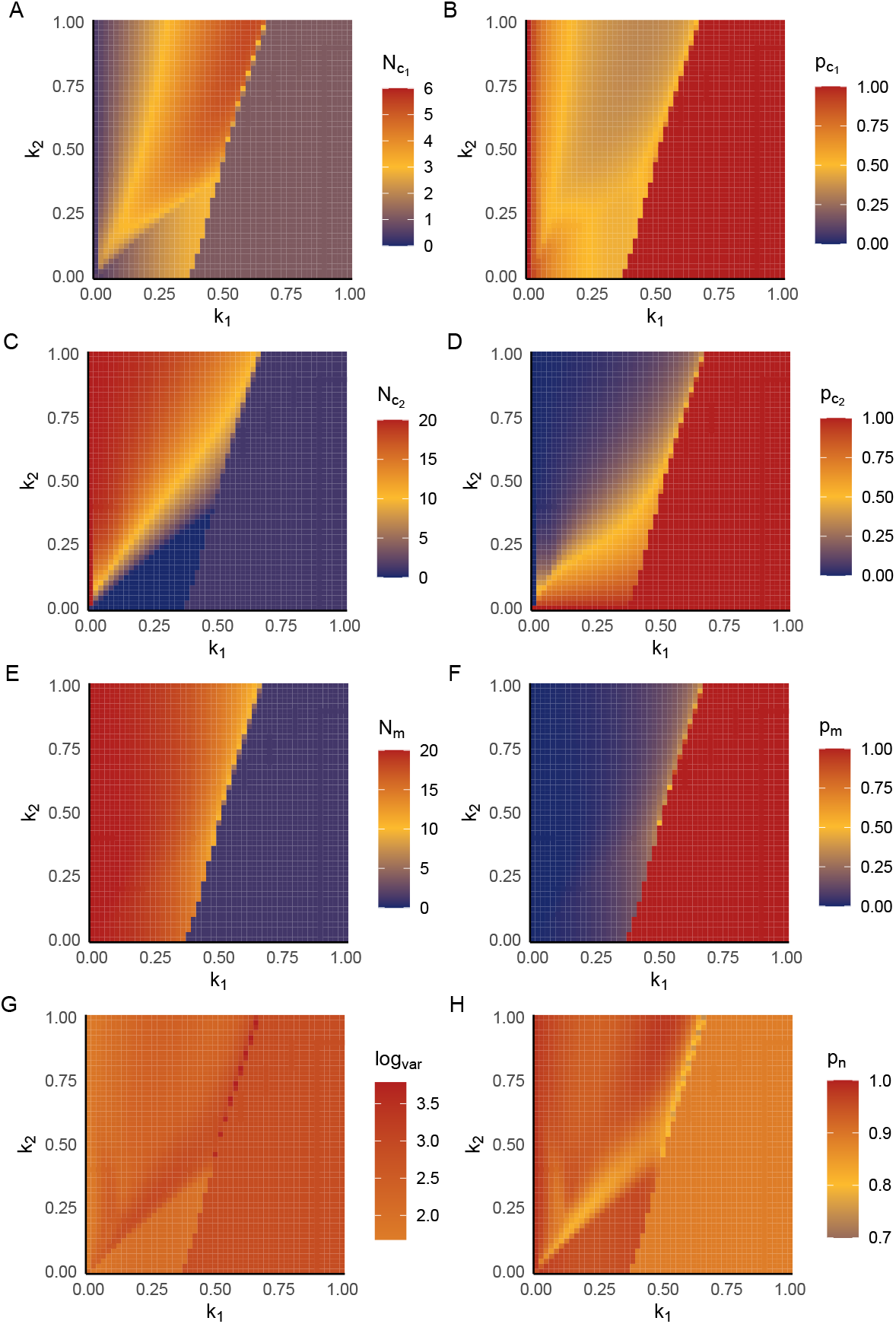
The effect of the Batesian mimic–model similarity (*k*_1_; x-axis) and the Müllerian mimic–model similarity (*k*_2_; y-axis) on the multi-mimicry system when recognition depends solely on morphological similarity (i.e., without abundance-dependent recognition). The different panels represent different variables: (A) Batesian mimic abundance 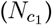, (B) attack probability on the Batesian mimic 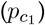, (C) the Müllerian mimic abundance 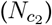, (D) attack probability on the Müllerian mimic 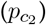, (E) model abundance (*N*_*m*_), (F) attack probability on the mimic (*p*_*m*_), (G) the log(variance) of the predator fluctuations, serving as an indicator of community stability, and (H) the attack probability on the alternative prey (*p*_*n*_). Note that the abundances of different state variables are on different scales to better show the pattern of each variable. See main text for other parameter values.

**Figure S5:**
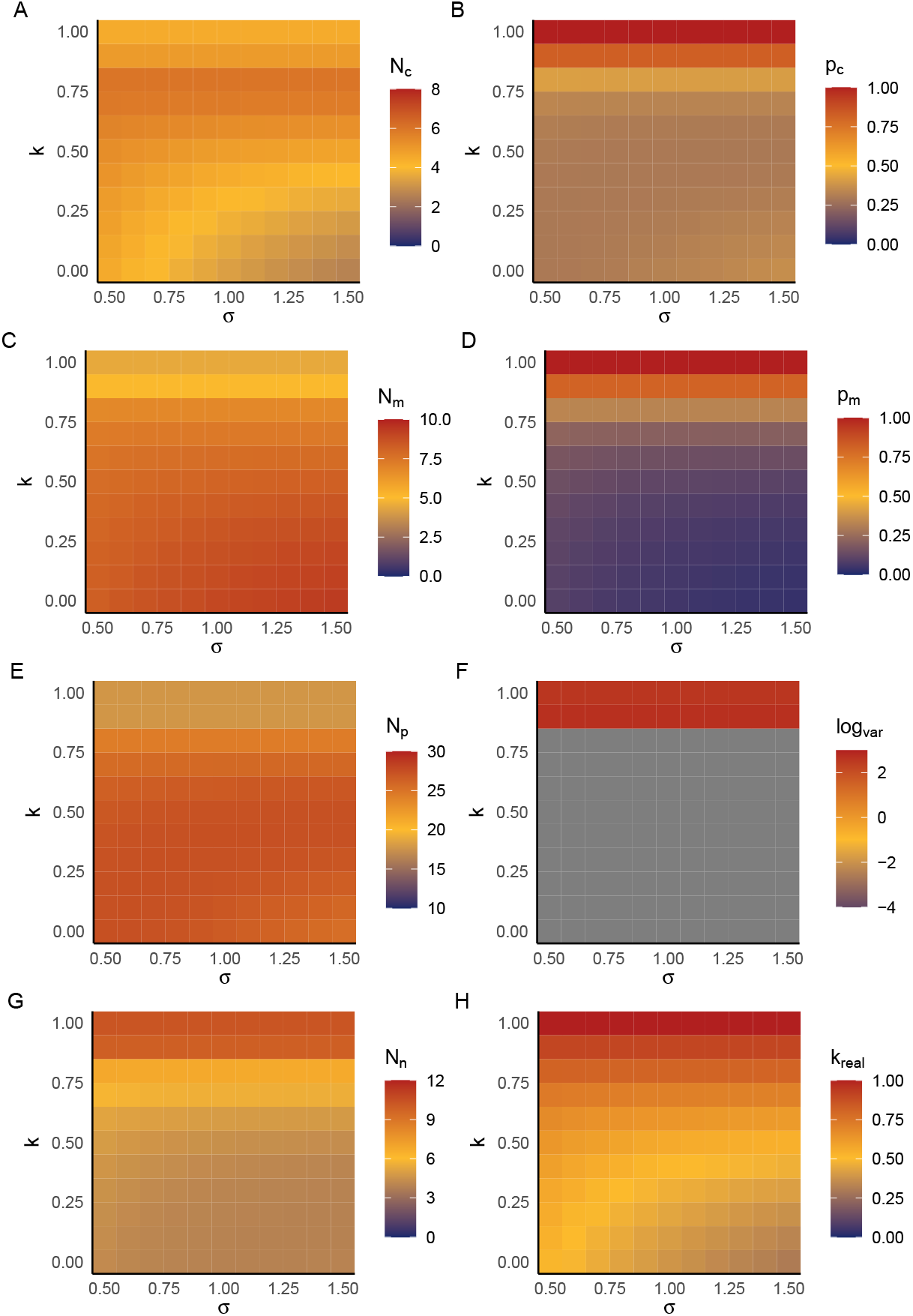
The effect of predator sensitivity to prey categories (*σ*; x-axis) and the mimic– model morphological similarity (*k*; y-axis) on the single Batesian mimicry system with abundance-dependent similarity. The different panels represent different variables: (A) Batesian mimic abundance (*N*_*c*_), (B) attack probability on the Batesian mimic (*p*_*c*_), (C) the model abundance (*N*_*m*_), (D) attack probability on the model (*p*_*m*_), (E) predator abundance (*N*_*p*_), (F) the log(variance) of the predator abundance fluctuation as an indicator of system stability, (G) alternative prey abundance (*N*_*n*_), and (H) the realized similarity due to abundance-dependent recognition (*p*_*n*_). Note that the abundances of different state variables are on different scales to better show the pattern of each variable. See main text for other parameter values.

